## Extended Data and SI for "Functional Integration of 3D-Printed Cerebral Cortical Tissue into a Brain Lesion"

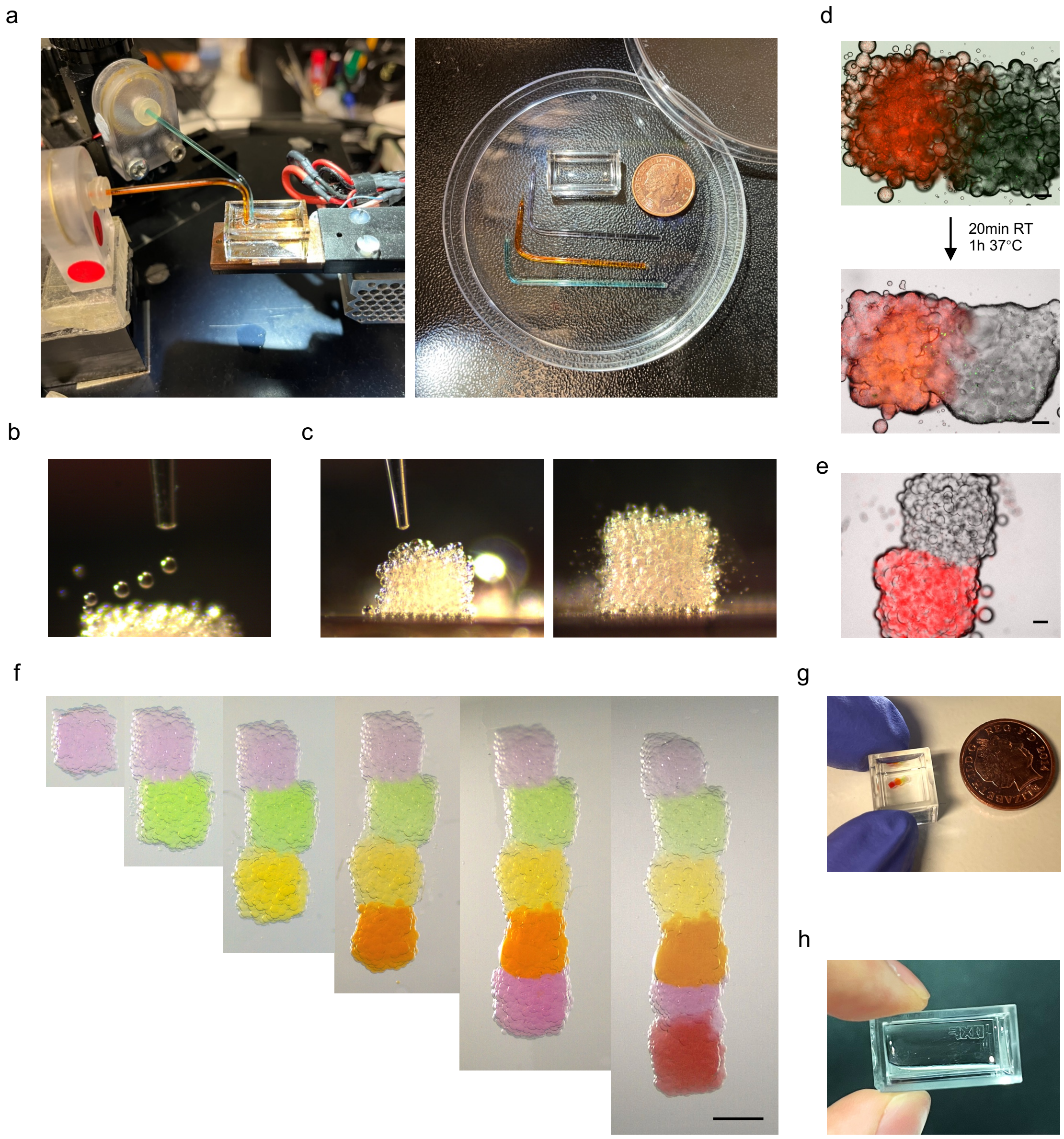


**Extended Data Fig. 1: Droplet-based 3D printing. a.** The droplet-based 3D bioprinter (left) and various components (right) including the glass printing cuvette and printing nozzles in comparison to a ten-pence coin. **b.** Side-view image of on-going printing. **c.** Side-views of printed droplet networks containing Matrigel only. **d.** Fluorescence images of a two-layered droplet network containing RFP-labelled UNPs and unlabelled DNPs. Raising the temperature, from room to physiological, facilitated gelation and annealing of printed two-layer networks. **e.** Fluorescence image of two-layered droplet network with fluorescent microbeads in one layer. **f.** Sequential generation of six-layered network by the droplets containing food dye coloured DPBS. Scale bar, 1000 µm. **g.** View of the six-layered network in **'f'**, in comparison to a ten-pence coin. **h.** View of the droplet network in Fig. **1l**. The network was printed as a mirror image of 'OXF' for imaging with an inverted microscope. For **'d'** and **'e'**: scale bar, 200 µm.

a

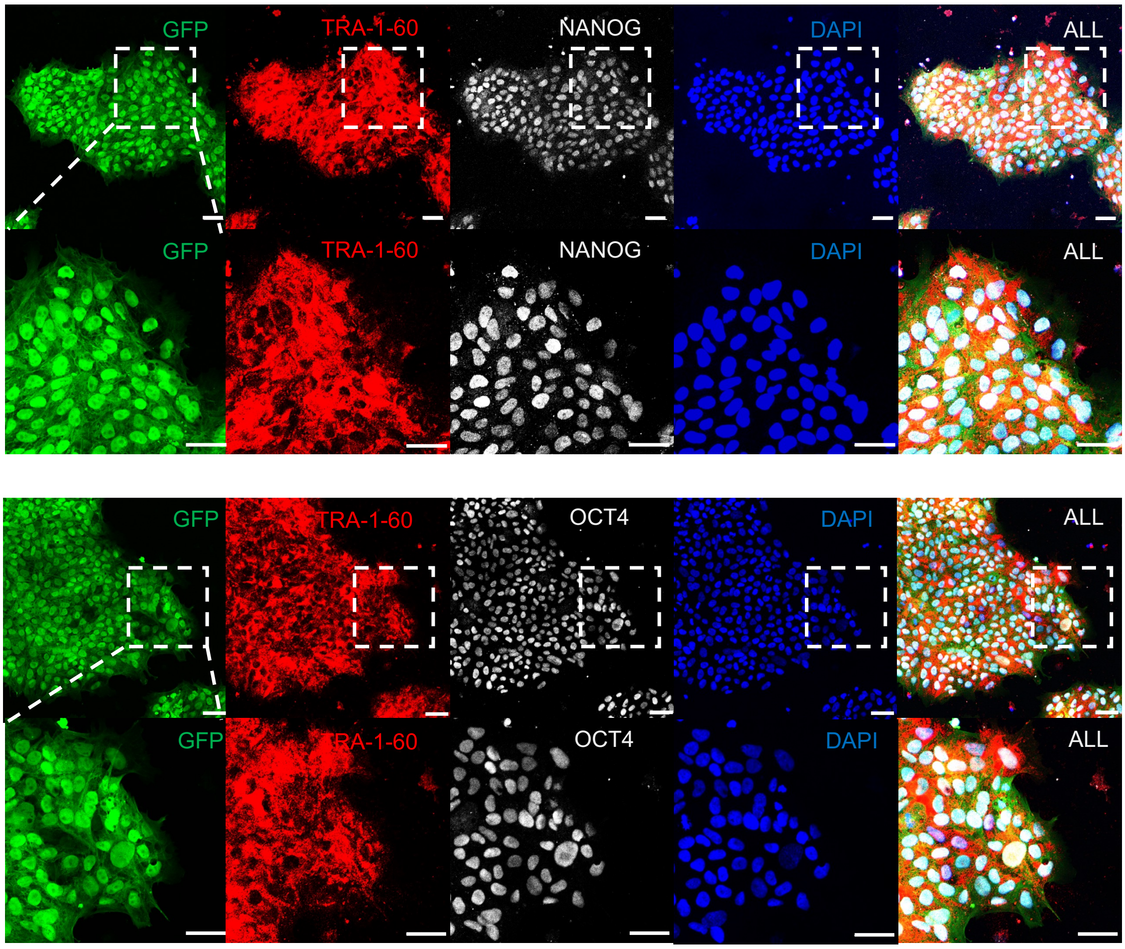


**Extended Data Fig. 2: Characterisation of human induced pluripotent stem cells (hiPSCs). a.** Confocal fluorescence images of immunostained hiPSCs showing the expression of pluripotent stem-cell markers TRA-1-60, NANOG and OCT4 in majority of the cells. Images at higher magnification of the regions indicated by the dashed boxes are shown in the second row. Scale bar, 50 μm.

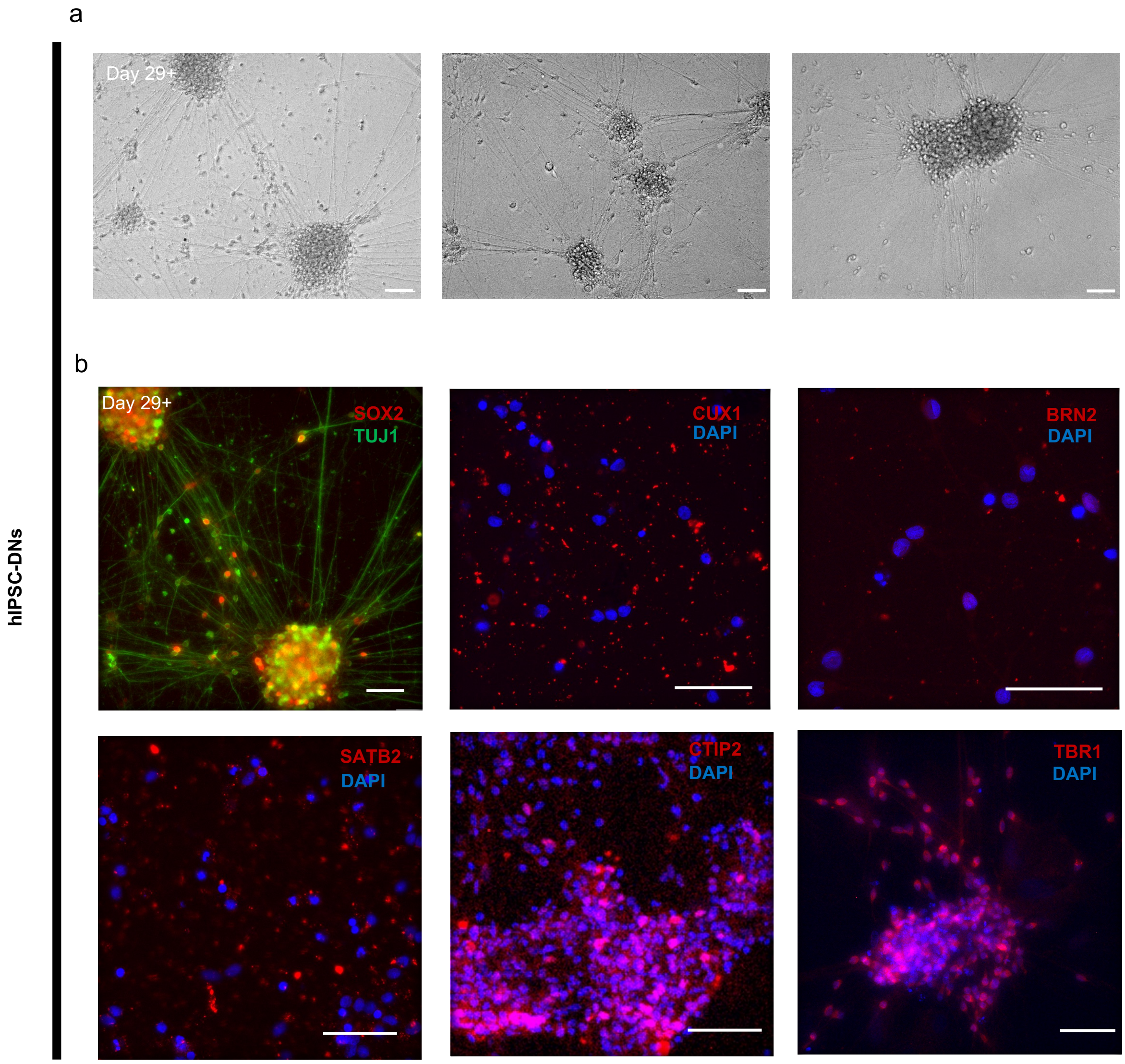


**Extended Data Fig. 3: Characterisation of hiPSCs derived deep-layer neurons (hiPSCs-DNs). a.** Bright-field images of DIV29+ hiPSCs-DNs showing mature neural morphology. **b.** Immunostaining of DNs showing expression of the neural stem cell marker SOX2, the general young neuron markers TUJ1 and the deep-layer markers (CTIP2 and TBR1). Expression of upper-layer markers (CUX1 and BRN2) and the middle-upper-layer marker (SATB2) are not detected despite high fluorescence intensity was used to reveal the background. For all panels: scale bar, 50 µm.

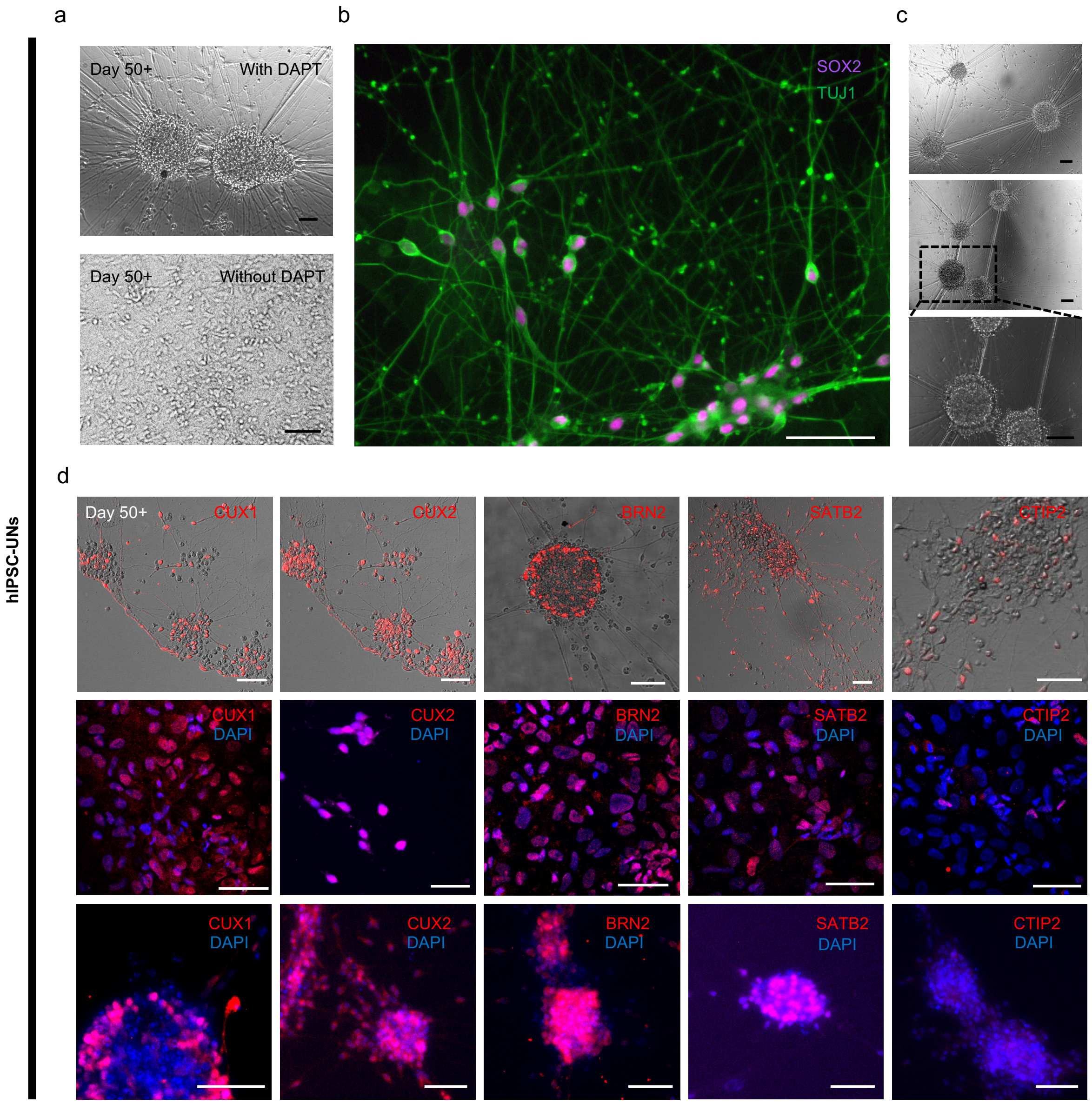


**Extended Data Fig. 4: Characterisation of hiPSC-derived upper-layer neurons (hiPSCs-UNs). a.** Bright-field images of DIV 50+ hiPSCs-UNs demonstrating the neuronal morphologies with (top) and without (bottom) DAPT treatment during maturation. UNs matured in NTM with DAPT are shown in **b**-**d**. **b.** DIV 50+ UNs immunostained with the neural stem cell marker SOX2 and the general young neuron marker TUJ1. **c.** Bright-field images hiPSCs-UNs showing mature morphology as indicated by extensive process outgrowth. **d.** Fluorescence images of DIV 50+ UNs from three independent experiments on each row showing expression of upper-layer markers (CUX1, CUX2 and BRN2) and the middle-upper-layer marker (SATB2), but low expression of the deep-layer marker (CTIP2). For all panels: scale bar, 50 μm.

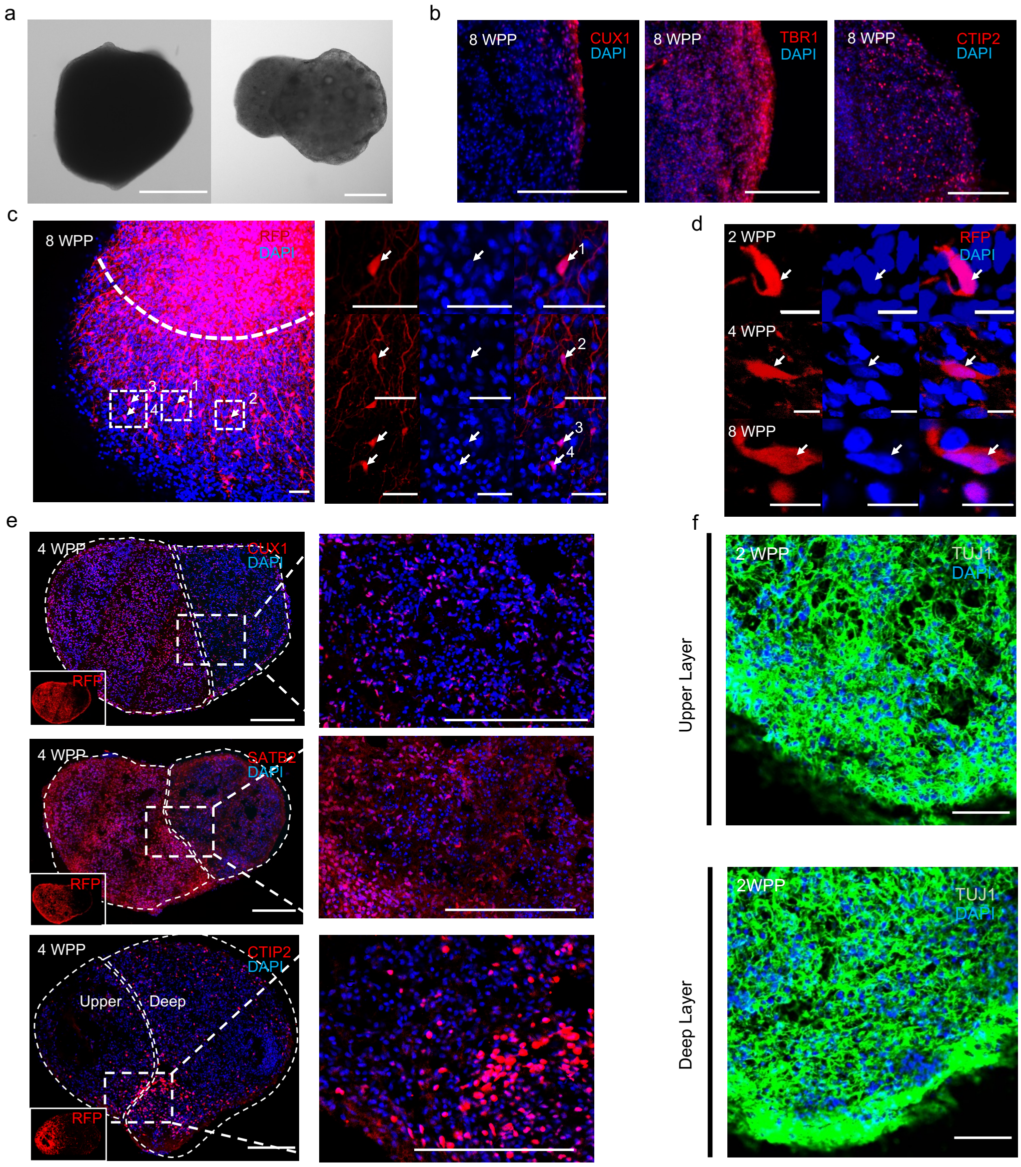


**Extended Data Fig. 5: Further characterisation of droplet-printed cerebral cortical tissues. a.** Bright-field images of one 2 WPP single-layer (with DNs,top) and one two-layer (with DNs and UNs, bottom) cortical tissues. **b.** Fluorescence images of sectioned 8 WPP deep-layer cortical tissues showing the abundant expression of the deep layer markers (CTIP2 & TBR1), and the sparse expression of upper-layer marker (CUX1). **c.** Confocal z-projection image (left) and high magnification images (right) showing cross-layer neuron migration in printed two-layer tissue at 8 WPP, visualized by RFP expression in UNs and DAPI nuclear staining in both UN and DNs. Dashed boxes indicate the magnified regions. Arrows and numbers indicate migrating neurons. Scale bar, 50 µm. **d.** Confocal images of 30 µm-thickness sections of 2, 4 and 8 WPP two-layer tissues showing cross-layer neuron migration, visualized by RFP and DAPI co-localisation. Scale bar, 10 µm. **e.** Immunofluorescence images at 4WPP of sectioned two-layer tissues showing expression of the layer-specific markers (CUX1, SATB2 and CTIP2). Bottom left small shows RFP expression of the tissue. Dashed lines outline the layers and dashed boxes indicate the magnified areas. **f.** Confocal images of sectioned two-layer tissues showing the expression of young neuronal marker TUJ1. Scale bar, 50 µm. For panels **'a'**, **'b'** & **'e'**: scale bar, 200 µm.

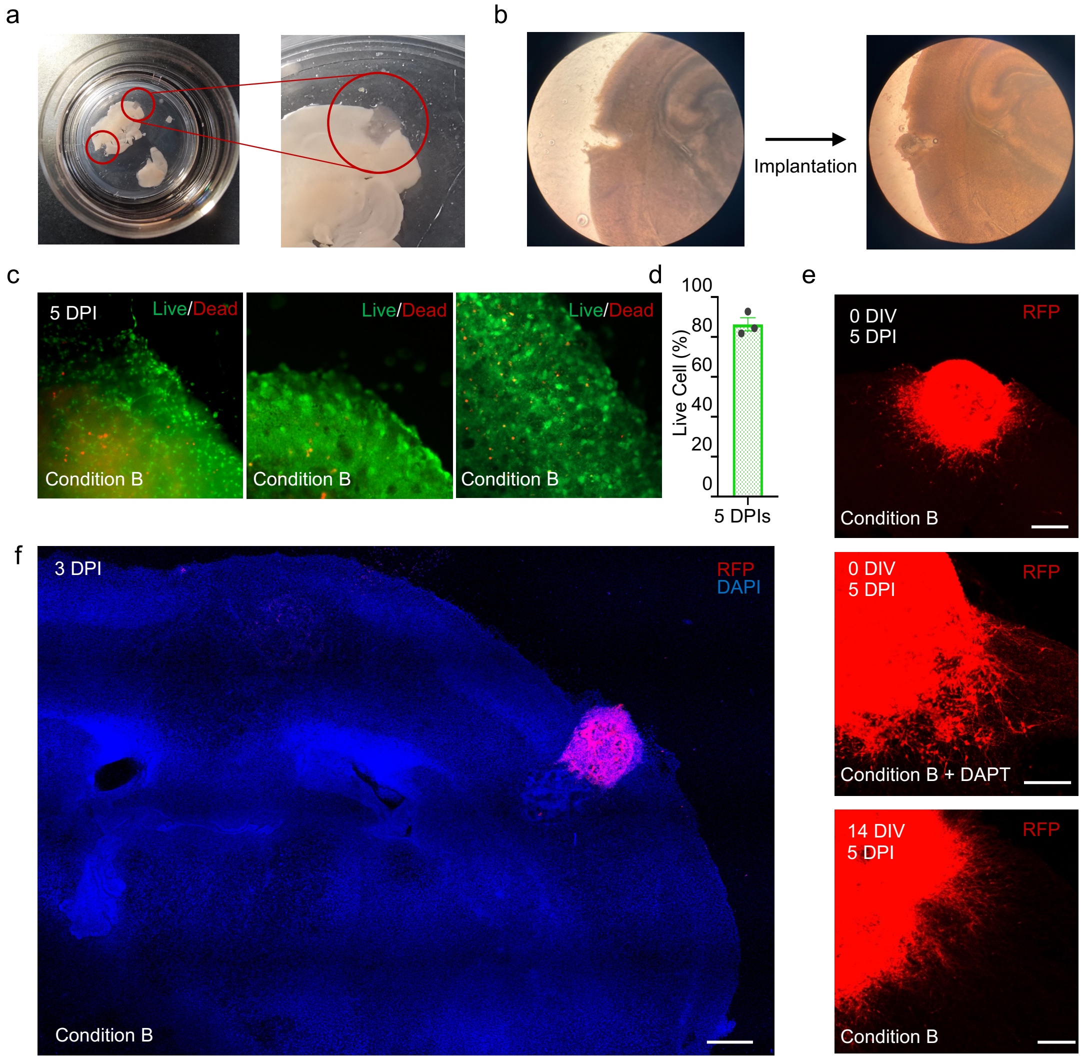


**Extended Data Fig. 6: Characterisation of implanted mouse brain explants. a.** 0 DPI explant with a lesion in the left cerebral hemisphere and a lesion implanted with printed deep-layer tissue in the right cerebral hemisphere. Right, a magnified image of lesion on the right hemisphere implanted with printed deep-layer cortical tissue. **b.** A bright-field image of a 0 DPI explant with a lesion implanted with a printed deep-layer cortical tissue. **c.** Fluorescence images of a live/dead assay of deep-layer cortical tissue implanted explant cultured under condition B at 5 DPIs. **d.** Quantitative live/dead analysis of host cells of 5 DPIs at condition B (n = 3). **e.** Further examples of implanted RFP-labelled deep-layer tissues under different nutrient conditions and with different pre-implantation culture times. **f.** Tiled fluorescence confocal image of an explant implanted with a two-layer printed tissue in the right hemisphere. Cells were visualized by RFP (UNs) and DAPI nuclear staining in UNs, DNs and the host. For all panels: scale bar, 200μm.

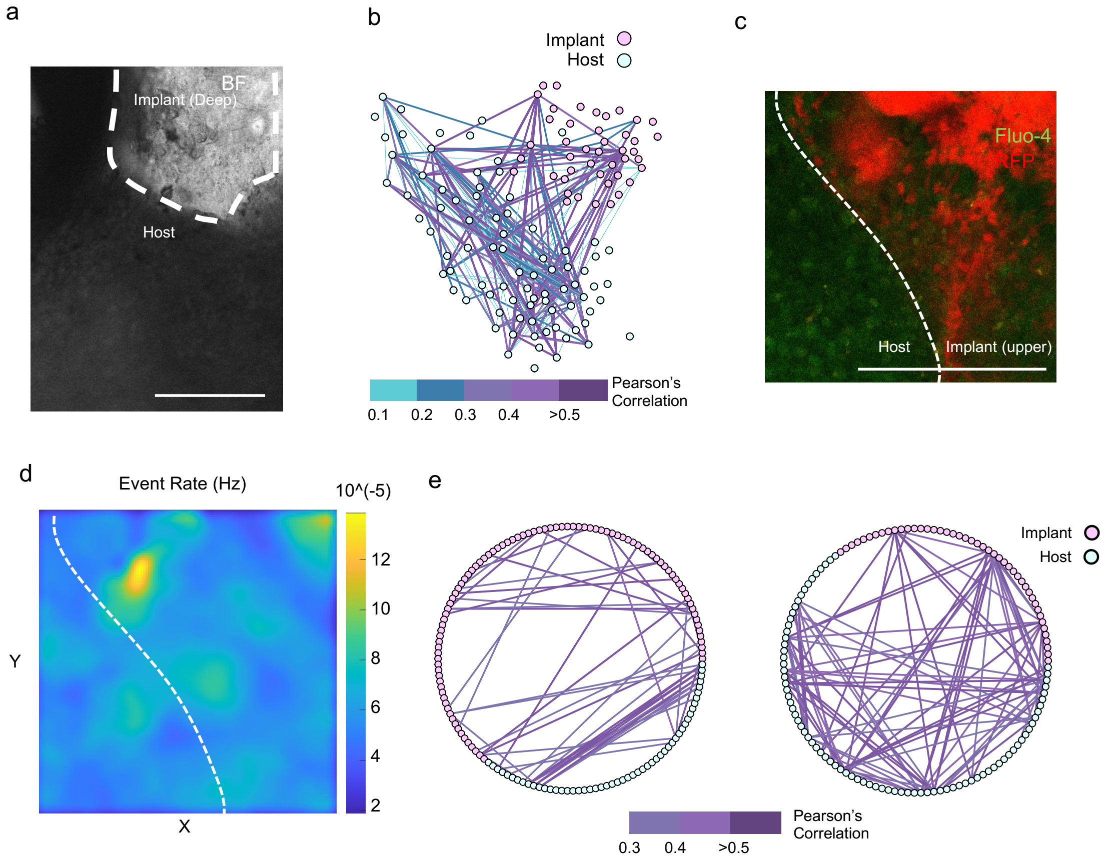


**Extended Data Fig. 7. Functional analysis of implanted mouse brain explants.** **a.** Bright-field image at 5 DPIs of an explant implanted with deep-layer cortical tissue (as indicated in **'Fig. 5h-j')**. The contrast difference between the implant and host marks the border between them. **b.** Network analysis of firing-correlated neurons between the implant and the host in **'Extended Data Fig.7a'** on 5 DPI. Circles correspond to neurons and lines indicate correlated firings. **c.** Fluorescence image of an explant implanted with upper-layer tissue, as also indicated on **'Fig. 5k-n'**. The RFP-labelled UNs and Fluo-4 labelled implant and host tissue mark the implant-host interface. **d.** Heatmap of neuron firing rate showing comparable neuron activity between explant and the implanted tissue. **e.** Network analysis of firing-correlated neurons for the 5 DPI implanted explants found in **'Fig.5h'** (Left) and **'Fig.5k'** (Right) in a circular layout. For all panels: scale bar, 200μm.

### Supplementary Information

**Functional Integration of 3D Printed Cerebral Cortical Tissue into a Brain Explant**

Yongcheng Jin[1], Ellina Mikhailova[1], Ming Lei[2], Sally Cowley[3], Tianyi Sun[2], Xingyun Yang[1], Yujia Zhang[1], Kaili Liu[4], Daniel Catarino[4], Luana Campos Soares[4], Sara Bandiera[4], Francis G. Szele[4]*, Zoltan Molnar[4]*, Linna Zhou[1,5]* and Hagan Bayley[1]*

[1]Department of Chemistry, University of Oxford, Oxford, OX1 3TA, United Kingdom.

[2]Department of Pharmacology, University of Oxford, Oxford, OX1 3QT, United Kingdom.

[3]James and Lillian Martin Centre for Stem Cell Research, Sir William Dunn School of Pathology, University of Oxford, South Parks Road, Oxford, OX1 3RE, United Kingdom.

[4]Department of Physiology, Anatomy and Genetics, University of Oxford, Oxford, OX1 3PT, United Kingdom.

[5]Ludwig Institute for Cancer Research, Nuffield Department of Medicine, University of Oxford, Oxford, OX3 7DQ, United Kingdom.

*

### Supplementary Table 1. Culture Medium Formula

| **Neural Induction Medium (NIM) 100mL** | | | | | |
|---|---|---|---|---|---|
| **Item** | **Volume** | **Final Conc** | **Stock Conc** | **Supplier** | **Cat no** |
| DMEM/F12 Medium | ~49 mL | NA | 1X | Life Technologies | 21331020 |
| Neurobasal Medium | ~49 mL | NA | 1X | Life Technologies | 21103-049 |
| B27 supplement | 1 mL | NA | NA | Life Technologies | 17504044 |
| N2 supplement | 0.5 mL | NA | NA | Life Technologies | 17502-048 |
| GlutaMax | 1 mL | NA | 100X | Life Technologies | 35050-038 |
| LDN193189 | 10 µL | 100nM | 1 mM | Sigma | SML0559 |
| SB431542 | 100 µL | 10 µM | 10 mM | Cambridge Bioscience | ZRD-SB-50 |
| Puromycin (opt) | 50 µL | 2.5µg/ml | 5 mg/ml | MP Biomedicals UK | 210055225 |
| **Neural Maintenance Medium (NMM) 100mL** | | | | | |
| **Item** | **volume** | **Final Conc** | **Stock Conc** | **Supplier** | **Cat no** |
| DMEM/F12 Medium | ~49 mL | NA | 1X | Life Technologies | 21331020 |
| Neurobasal Medium | ~49 mL | NA | 1X | Life Technologies | 21103-049 |
| B27 supplement | 1 mL | NA | NA | Life Technologies | 17504044 |
| N2 supplement | 0.5 mL | NA | NA | Life Technologies | 17502-048 |
| GlutaMax | 1 mL | NA | 100X | Life Technologies | 35050-038 |
| Puromycin (opt) | 50 µL | 2.5µg/ml | 5 mg/ml | MP Biomedicals UK | 210055225 |
| **Neural Maintenance Medium + Growth Factors (NMM + GFs) 100mL** | | | | | |
| **Item** | **volume** | **Final Conc** | **Stock Conc** | **Supplier** | **Cat no** |
| DMEM/F12 Medium | ~49 mL | NA | 1X | Life Technologies | 21331020 |
| Neurobasal Medium | ~49 mL | NA | 1X | Life Technologies | 21103-049 |
| B27 supplement | 1 mL | NA | NA | Life Technologies | 17504044 |
| N2 supplement | 0.5 mL | NA | NA | Life Technologies | 17502-048 |
| GlutaMax | 1 mL | NA | 100X | Life Technologies | 35050-038 |
| Fibroblast Growth Factor-2 (FGF-2) | 10 µL | 10 ng/mL | 100µg/mL | R&D Systems | 4114-TC-01M |
| Epidermal Growth Factor (EGF) | 10 µL | 10 ng/mL | 100µg/mL | Life Technologies | PHG0311 |
| Brain-derived Neurotrophic Factor (BDNF) | 10 µL | 10 ng/mL | 100µg/mL | Life Technologies | PHC7074 |
| **Freezing Medium 10 mL** | | | | | |
| **Item** | **volume** | **Final Conc** | **Stock Conc** | **Supplier** | **Cat no** |
| ESC-qualified FBS | 9 mL | NA | NA | Gibco | 16141061 |
| DMSO | 1 mL | NA | NA | Merck | D2650-100ML |
| **Neural Terminal Medium (NTM) 100 mL** | | | | | |
| **Item** | **volume** | **Final Conc** | **Stock Conc** | **Supplier** | **Cat no** |
| Neurobasal Medium | 97 mL | NA | 1X | Life Technologies | 21103-049 |
| B27 supplement | 2 mL | NA | NA | Life Technologies | 17504044 |
| GlutaMax | 1 mL | NA | 100X | Life Technologies | 35050-038 |
| DAPT | 10 µL | 10 µM | 100 mM | Tocris | Oct-34 |
| Puromycin (opt) | 50 µL | 2.5µg/ml | 5 mg/ml | MP Biomedicals UK | 210055225 |
| **Brain Explant Culture Medium Condition A 100mL** | | | | | |
| **Item** | **volume** | **Final Conc** | **Stock Conc** | **Supplier** | **Cat no** |
| DMEM/F12 Medium | ~36 mL | NA | 1X | Life Technologies | 21331020 |
| Neurobasal Medium | ~36 mL | NA | 1X | Life Technologies | 21103-049 |
| B27 supplement | 1 mL | NA | NA | Life Technologies | 17504044 |
| N2 supplement | 0.5 mL | NA | NA | Life Technologies | 17502-048 |
| GlutaMax | 1 mL | NA | 100X | Life Technologies | 35050-038 |
| Pen/Strep | 1 mL | NA | 10,000 U/mL | Gibco | 15140122 |
| Horse Serum | 25 mL | NA | NA | Life Technologies | 16050130 |
| **Brain Explant Medium Condition B 100mL** | | | | | |
| **Item** | **volume** | **Final Conc** | **Stock Conc** | **Supplier** | **Cat no** |
| Brainphys Neural Medium | 72.5 mL | NA | NA | Stemcell Technologies | 5792 |
| SM1 supplement | 1.5 mL | NA | NA | Stemcell Technologies | 5792 |
| Pen/Strep | 1 mL | NA | 10,000 U/mL | Gibco | 15140122 |
| Horse Serum | 25 mL | NA | NA | Life Technologies | 16050130 |

### Supplementary Table 2. Consumables

| Consumables (Cell Culture) | | | | | |
|---|---|---|---|---|---|
| **Item** | **volume** | **Final Conc** | **Stock Conc** | **Supplier** | **Cat no** |
| mTeSR Plus Medium | NA | NA | NA | Stemcell Technologies | 100-0276 |
| Geltrex | NA | NA | NA | Gibco | A1413302 |
| StemPro Accutase | NA | NA | NA | Life Technologies | A1110501 |
| DPBS | NA | NA | NA | Gibco | 14190144 |
| UltraPure 0.5M EDTA | NA | 0.5 mM | 0.5 M | Life Technologies | 15575020 |
| Distilled Water | NA | NA | NA | Life Technologies | 15230089 |
| Y-27632 | NA | 10 µM | 1mM | Abcam | ab120129-10mg |
| ReLeSR | NA | NA | NA | Stemcell Technologies | 5872 |
| **Consumables (Droplet Printing)** | | | | | |
| **Item** | **volume** | **Final Conc** | **Stock Conc** | **Supplier** | **Cat no** |
| Silicone oil AR20 | NA | NA | NA | Sigma | 10836 |
| Undecane | NA | NA | NA | Sigma | 1120-21-4 |
| DPHPC | NA | NA | NA | Avanti | 850356 |
| Trimethoxysilane | NA | 5% v/v | NA | Sigma | 281778 |
| Matrigel | NA | NA | NA | Corning | 354230 |
| **Consumables (Brain Explant)** | | | | | |
| **Item** | **volume** | **Final Conc** | **Stock Conc** | **Supplier** | **Cat no** |
| EBSS | NA | NA | NA | Life Technologies | 24010043 |
| Culture Insert | NA | NA | NA | Merck | PICM0RG50 |
| X30 Cell Imaging Dish, | NA | NA | NA | Fisher Scientific UK | 15670537 |
| BrainPhys Imaging Optimized Medium | NA | NA | NA | Stemcell Technologies | 5796 |
| UltraPure Low Melting Point Agarose | NA | NA | NA | Life Technologies | 16520050 |
| Microtome blade | NA | NA | NA | Fisher Scientific | 11912355 |
| Needle | NA | NA | NA | Fisher Scientific | 10749891 |
| Super Glue | NA | NA | NA | Office Depot | 4086446 |
| **Consumables (qPCR)** | | | | | |
| **Item** | **volume** | **Final Conc** | **Stock Conc** | **Supplier** | **Cat no** |
| LunaScript(R) RT SuperMix Kit | NA | NA | NA | New England Biolabs | E3010L |
| Monarch(R) Total RNA Miniprep Kit | NA | NA | NA | New England Biolabs | T2010S |
| Luna(R) Universal qPCR Master Mix | NA | NA | NA | New England Biolabs | M3003L |
| MicroAmp Fast Optical 96-Well Reaction Plate | NA | NA | NA | Life Technologies | 4346906 |
| Nuclease-Free water | NA | NA | NA | QIAGEN | 129114 |
| **Consumables (Immunostaining, live/dead assay and Fluo-4 imaging)** | | | | | |
| **Item** | **volume** | **Final Conc** | **Stock Conc** | **Supplier** | **Cat no** |
| Ibidi µ-Slide 18 Well | NA | NA | NA | ThistleScientific | SKU 81816 |
| Paraformaldehyde 4% | NA | 4% | 4% | Alfa Aesar | J61899.AK |
| Glycine 1 M Solution | NA | NA | NA | Merck | 67419-1ML-F |
| Triton X-100 | NA | NA | NA | Merck | 93443-100ML |
| Tween-20 | NA | NA | NA | Alfa Aesar | P9416-50ML |
| Normal Goat Serum | NA | NA | NA | Abcam | ab7481 |
| Normal Donkey Serum | NA | NA | NA | Abcam | ab7475 |
| DAPI Solution | NA | 1X | 10000X | Merck | MBD0015-1ML |
| Mounting Medium With DAPI | NA | 1X | 1X | Abcam | ab104139 |
| Fluo-4 Direct Calcium Assay Kit | NA | 1X | 2X | Life Technologies | F10471 |
| Calcein-AM | NA | 2.5 µM | NA | Cambridge bioscience | 1755-50 |
| Propidium iodide | NA | 5.0 µM | NA | Sigma | P4170 |
| **Plastics and others** | | | | | |
| 1.8ml Cryogenic Vial | NA | NA | NA | STARLAB | E3090-6222 |
| 10µl Pipette Tip | NA | NA | NA | STARLAB | S1121-2710 |
| 20µl Pipette Tip | NA | NA | NA | STARLAB | S1120-1710 |
| 200µl Pipette Tip | NA | NA | NA | STARLAB | S1126-7810 |
| 1000µl Pipette Tip | NA | NA | NA | STARLAB | S1120-8810 |
| 6 Well Tissue Culture Plate | NA | NA | NA | Greiner Bio-One | 657160 |
| 12 Well Tissue Culture Plate | NA | NA | NA | Greiner Bio-One | 665180 |
| 24Well Tissue Culture Plate | NA | NA | NA | Greiner Bio-One | 662160 |
| 48 Well Tissue Culture Plate | NA | NA | NA | Greiner Bio-One | 677180 |
| 96 well Tissue Culture Plate | NA | NA | NA | Greiner Bio-One | 655180 |
| 96 well Assessment Plate | NA | NA | NA | Corning | CLS3603 |
| 5mL Stripette | NA | NA | NA | Scientific Laboratory Supp | 4487 |
| 10mL Stripette | NA | NA | NA | Scientific Laboratory Supp | 4488 |
| 25mL Stripette | NA | NA | NA | Scientific Laboratory Supp | 4489 |
| Cryo Container | NA | NA | NA | VWR International | 479-3200 |
| Microslides | NA | NA | NA | VWR International | 631-0448 |
| PAP Pen | NA | NA | NA | Merck | Z672548-1EA |

### Supplementary Table 3. Antibodies and Primers

**Primary Antibodies**

| Target | Original Species | Manufacturer | Dilution Factor | Cat. No |
|---|---|---|---|---|
| CUX1 | Ms | Santa Cruz Biotechnology | 100 | sc-13024 |
| CUX2 | Rb | AbCam | 200 | ab216588 |
| BRN2 | Ms | Santa Cruz Biotechnology | 100 | sc-393324 |
| SATB2 | Rb | AbCam | 200 | ab92446 |
| CTIP2 | Rat | AbCam | 200 | ab18465 |
| TBR1 | Rb | Merck | 500 | AB10554 |
| SOX2 | Rb | Millipore | 100-200 | ab5603 |
| TUJ1 | Ms | AbCam | 500-1000 | ab78078 |
| GFAP | Rat | Invitrogen | 200 | 13-0300 |
| HNCAM | Rb | AbCam | 200 | ab75813 |

**Secondary Antibodies**

| Target Species | Original Species | Fluorophore | Manufacturer | Dilution Factor | Cat. No |
|---|---|---|---|---|---|
| Rb | Goat | Alex488 | Invitrogen | 1000 | a11006 |
| Ms | Goat | Alex488 | Invitrogen | 1000 | a32723 |
| Ms | Goat | Alex633 | Invitrogen | 1000 | a21052 |
| Rat | Goat | Alex647 | Invitrogen | 1000 | a21247 |
| Rb | Goat | Alex647 | Invitrogen | 1000 | a21245 |

**qPCR Primer**

| Target | Forward or Backward | 5'-3' Sequence | Manufacturer |
|---|---|---|---|
| PAX6 | F | GCCAGCAACACACCTAGTCA | Life Technologies |
| | R | TGTGAGGGCTGTGTCTGTTC | Life Technologies |
| Nestin | F | GGAAGAGAACCTGGGAAAGG | Life Technologies |
| | R | CTTGGTCCTTCTCCACCGTA | Life Technologies |
| CTIP2 | F | GAGTACTGCGGCAAGGTGTT | Life Technologies |
| | R | TAGTTGCACAGCTCGCACTT | Life Technologies |
| BRN2 | F | GACCTTTGCAGGCGAGTAAC | Life Technologies |
| | R | TCAGGAAGCTGCATTTTGTG | Life Technologies |
| CUX1 | F | GCTCTCATCGGCCAATCACT | Life Technologies |
| | R | TCTATGGCCTGCTCCACGT | Life Technologies |
| CUX2 | F | AAGGAGATCGAGTCGCAGAA | Life Technologies |
| | R | CTCCAGGATGCTCTTGATGG | Life Technologies |
| 18S | F | GAGGATGAGGTGGAACGTGT | Life Technologies |
| | R | TCTTCAGTCGCTCCAGGTCT | Life Technologies |

### Supplementary Table 4. Detailed Statistical Test

| Figure | Comparison | P value | Catelogue | Label | Test |
|---|---|---|---|---|---|
| Fig. 2d | CUX1 vs CTIP2 | 0.0009 | P<0.001 | *** | One-way ANOVA with Dunnett's test |
| | BRN2 vs CTIP2 | 0.0004 | P<0.001 | *** | One-way ANOVA with Dunnett's test |
| | SATB2 vs CTIP2 | 0.0008 | P<0.001 | *** | One-way ANOVA with Dunnett's test |
| Fig. 2e | DNPs CUX1 vs UNPs CUX1 | 0.022 | P<0.05 | * | Unpaired Student t-test |
| | DNPs CUX1 vs UNs CUX1 | 0.0023 | P<0.01 | ** | Unpaired Student t-test |
| | UNPs CUX1 vs UNs UCX1 | 0.7258 | P>0.05 | ns | Unpaired Student t-test |
| Fig. 3g | RFP Coverage Upper 4WPP vs 2WPP | 0.4265 | P>0.05 | ns | One-way ANOVA with Dunnett's test |
| | RFP Coverage Upper 8WPP vs 2WPP | 0.1339 | P>0.05 | ns | One-way ANOVA with Dunnett's test |
| | RFP Coverage Deep 4WPP vs 2WPP | 0.0599 | P>0.05 | ns | One-way ANOVA with Dunnett's test |
| | RFP Coverage Deep 8WPP vs 2WPP | 0.0115 | P<0.05 | * | One-way ANOVA with Dunnett's test |
| | RFP Coverage D/U 4WPP vs 2WPP | 0.0074 | P<0.01 | ** | One-way ANOVA with Dunnett's test |
| | RFP Coverage D/U 8WPP vs 2WPP | 0.0003 | P<0.001 | *** | One-way ANOVA with Dunnett's test |
| | Migration 4WPP vs 2WPP | 0.463 | P>0.05 | ns | One-way ANOVA with Dunnett's test |
| | Migration 8wpp VS 2WPP | 0.0093 | P<0.01 | ** | One-way ANOVA with Dunnett's test |
| Fig. 3i | 2WPP-CUX1 Upper vs Deep | 0.0042 | P<0.01 | ** | Unpaired Student t-test |
| | 4WPP-CUX1 Upper vs Deep | 0.0032 | P<0.01 | ** | Unpaired Student t-test |
| | 8WPP-CUX1 Upper vs Deep | 0.002 | P<0.01 | ** | Unpaired Student t-test |
| | 2WPP-SATB2 Upper vs Deep | 0.0403 | P<0.05 | * | Unpaired Student t-test |
| | 4WPP-SATB2 Upper vs Deep | 0.0477 | P<0.05 | * | Unpaired Student t-test |
| | 8WPP-SATB2 Upper vs Deep | 0.027 | P<0.05 | * | Unpaired Student t-test |
| | 2WPP-CTIP2 Upper vs Deep | 0.0045 | P<0.01 | ** | Unpaired Student t-test |
| | 4WPP-CTIP2 Upper vs Deep | 0.0553 | P>0.05 | ns | Unpaired Student t-test |
| | 8WPP-CTIP2 Upper vs Deep | 0.3324 | P>0.05 | ns | Unpaired Student t-test |
| | 2WPP-TUJ1 Upper vs Deep | 0.2852 | P>0.05 | ns | Unpaired Student t-test |
| | 4WPP-TUJ1 Upper vs Deep | 0.6063 | P>0.05 | ns | Unpaired Student t-test |
| | 8WPP-TUJ1 Upper vs Deep | 0.6958 | P>0.05 | ns | Unpaired Student t-test |
| Fig. 4g | ConditionB -DAPT vs Condition A -DAPT | 0.015 | P<0.05 | * | Unpaired Student t-test |
| | ConditionB +DAPT vs Condition B -DAPT | 0.0077 | P<0.01 | ** | Unpaired Student t-test |
| | Effect of ConditionA vs ConditionB | 0.0008 | P<0.001 | *** | Two-way ANOVA |
| | Effect of +DAPT vs -DAPT | 0.0014 | P<0.01 | ** | Two-way ANOVA |
| Fig. 4i | 14 Days vs 0 Days | 0.0211 | P<0.05 | * | Unpaired Student t-test |
| Fig. 4k | UNs 1DPIs vs UNs 3DPIs | 0.0487 | P<0.05 | * | One-way ANOVA with Dunnett's test |
| | UNs 1DPIs vs UNs 5DPIs | 0.0015 | P<0.01 | ** | One-way ANOVA with Dunnett's test |
| | UNs 5DPIs vs 1d DNs 5DPIs | 0.4676 | P>0.05 | ns | Unpaired Student t-test |
| Fig. 5b | 1DPI Upper vs Deep | 0.2045 | P>0.05 | ns | Unpaired Student t-test |
| | 3DPI Upper vs Deep | 0.204 | P>0.05 | ns | Unpaired Student t-test |
| | 5DPI Upper vs Deep | 0.1179 | P>0.05 | ns | Unpaired Student t-test |
| | 3 DPI Deep vs 1 DPI Deep | 0.0005 | P<0.001 | *** | Unpaired Student t-test |
| | 3 DPI Upper vs 1 DPI Upper | 0.0018 | P<0.01 | ** | Unpaired Student t-test |
| | 5 DPI Deep vs 1 DPI Deep | <0.0001 | P<0.0001 | **** | Unpaired Student t-test |
| | 5 DPI Upper vs 1 DPI Upper | <0.0001 | P<0.0001 | **** | Unpaired Student t-test |
| Fig. 5e | RFP Upper vs Host | <0.0001 | P<0.0001 | **** | One-way ANOVA with Dunnett's test |
| | RFP Deep vs Host | 0.9481 | P>0.05 | ns | One-way ANOVA with Dunnett's test |
| | HNCAM Upper vs Host | 0.0301 | P<0.05 | * | One-way ANOVA with Dunnett's test |
| | HNCAM Deep vs Host | 0.117 | P>0.05 | ns | One-way ANOVA with Dunnett's test |

**Supplementary Video 1**, 3D reconstructed confocal z-projection image showing cross-layer process outgrowth and neuron migration in a printed two-layer tissue at 8 WPP, visualized by RFP (false coloured as fire) expression in UNs and DAPI nucleus staining in both UN and DNs. Scale bars: 500 μm.

**Supplementary Video 2**, Fluo-4 calcium ion activity recording of the explant implanted with DNPs only at 5 DPI, as indicated in **'Fig. 5h'.** Scale bars: 200 μm.
